# Allogeneic CAR-invariant Natural Killer T Cells Exert Potent Antitumor Effects Through Host CD8 T Cell Cross-Priming

**DOI:** 10.1101/2021.02.03.428987

**Authors:** Federico Simonetta, Juliane K. Lohmeyer, Toshihito Hirai, Kristina Maas-Bauer, Maite Alvarez, Arielle S. Wenokur, Jeanette Baker, Amin Aalipour, Xuhuai Ji, Samuel Haile, Crystal L. Mackall, Robert S. Negrin

## Abstract

The development of allogeneic chimeric antigen receptor (CAR) T cell therapies for off-the-shelf use is a major goal yet faces two main immunological challenges, namely the risk of graft-versus-host-disease (GvHD) induction by the transferred cells and the rejection by the host immune system limiting their persistence. We demonstrate that allogeneic CAR-engineered invariant natural killer T (iNKT) cells, a cell population without GvHD-induction potential that displays immunomodulatory properties, exerted potent direct and indirect antitumor activity in murine models of B-cell lymphoma when administered across major MHC-barriers. In addition to their known direct cytotoxic effect, allogeneic CAR iNKT cells induced tumor-specific antitumor immunity through host CD8 T cell cross-priming, resulting in a potent antitumor effect lasting longer than the physical persistence of the allogeneic cells. The utilization of off-the-shelf allogeneic CAR iNKT cells could meet significant unmet needs in the clinic.

## Introduction

Chimeric antigen receptor (CAR) T cells have resulted in dramatic and effective therapy for a range of relapsed and refractory malignancies. The use of autologous cells for the generation of CAR T cells represents a significant limitation to their widespread use for a number of significant reasons, including the impact of disease and treatment on the T cell product, costs of individual production, and the time required to produce the cellular product for patients with often rapidly progressive disease. The development of universal allogeneic CAR T cells could address these challenges yet faces two major limitations, namely the risk of graft-versus-host-disease (GvHD) induction by the allogeneic cells that recognize host tissues and the rejection of the CAR modified cells by the host immune system. Ablation of the T-cell receptor (TCR) (1–4) or use of non-MHC-restricted innate lymphocytes have been attempted to prevent GvHD. Similarly, ablation of MHC-class-I molecules to limit rejection by the host immune-system (5) has been employed in preclinical models.

Invariant Natural Killer T (iNKT) cells are a rare subset of innate lymphocytes representing less then 1% of the total lymphocyte population both in humans and mice. iNKT cells express a semi-invariant TCR recognizing glycolipids presented in the context of the monomorphic, MHC-like molecule CD1d. Because of their peculiar TCR constitution and antigen recognition modality, iNKT cells do not display any GvHD induction potential and can even prevent GvHD (reviewed in (6)). iNKT cells display potent direct antitumor activity through production of cytotoxic molecules(7). Several groups successfully generated human CAR iNKT cells provided with antitumor potential as assessed *in vitro* and in xenogeneic murine models(8–12). These studies revealed several advantages of using CAR iNKT cells over conventional CAR T cells, including their lack of induction of xeno-GvHD(8), their preferential migration to tumor sites (8) and their capacity of CAR iNKT to target both the natural ligand CD1d and the CAR-targeted antigen (11). In addition to their direct cytotoxic effect, iNKT cells are known for their strong immunomodulatory effect. In particular, iNKT cells induce CD8 T cell cross-priming(13, 14) through the licensing of CD103+ CD8alpha dendritic cells (15–17) allowing the establishment of long-lasting antitumor CD8 T cell responses in murine models (18–21).

In this study, we tested the hypothesis that induction of host-CD8 T cell cross priming by allogeneic CAR iNKT cells would allow the establishment of an antitumor immunity lasting beyond the physical persistence of the transferred cells. Taking advantage of the immunoadjuvant role of iNKT cells and their lack of GvHD-inducing potential, we demonstrate that allogeneic CAR iNKT cells exert, in addition of their previously reported direct antitumor effect (8–12), an indirect effect through the induction of host CD8 T cell cross-priming.

## Materials and Methods

### Mice

BALB/cJ (H-2K^d^) and FVB/NJ (H-2K^q^) mice were purchased from the Jackson Laboratory (Sacramento, CA). Firefly Luciferase *(Luc+)* transgenic FVB/N mice have been previously reported (22) and were bred in our animal facility at Stanford University. BALB/c Rag1^-/-^ gamma-chain^-/-^ and BALB/c BATF3^-/-^ mouse strains were kind gifts of Dr. Irving Weissman and Dr. Samuel Strober respectively and were bred in our animal facility at Stanford University. All procedures performed on animals were approved by Stanford University’s Institutional Animal Care and Use Committee and were in compliance with the guidelines of humane care of laboratory animals.

### CAR iNKT and conventional CAR T generation

Murine CD19.28z CAR iNKT and conventional CAR T cells specifically recognizing the murine CD19 molecule were generated using an adaptation of previously reported protocols (23). Murine CD19 (mCD19) CAR stable producer cell line (24) was kindly provided by Dr. Terry J. Fry. iNKT cells were negatively enriched from FVB/N mouse spleen single-cell suspensions and using a mixture of biotinylated monoclonal antibodies (GR-1, clone: RB6-8C5; CD8a, clone: 53-6.7; CD19, clone: 6D5; TCRγδ, clone: GL3; TER119/erythroid cell, clone: TER-119; CD62L, clone: MEL-14; BioLegend) and negative selection by anti-biotin microbeads (BD IMag™ Streptavidin Particles Plus DM, BD Biosciences). The enriched fraction (typically 10-30% enrichment) was then stimulated for 5 days with a synthetic analog of α-galactosylceramide (KRN7000, 100 ng/ml, REGiMMUNE) in the presence of human IL-2 (100 UI/ml; NCI Repository) and human IL-15 (100 ng/ml; NCI Repository). Cells were grown in DMEM media supplemented with 10% heat-inactivated fetal bovine serum (FBS), 1 mmol/L sodium pyruvate, 2 mmol/L glutamine, 0.1 mmol/L nonessential amino acids, 100 U/mL penicillin, and 100 μg/mL streptomycin at 37°C with 5% CO_2_. Conventional T cells were enriched from FVB/N mouse spleen single-cell suspensions using the mouse Pan T Cell Isolation Kit II (Miltenyi Biotec) according to the manufacturer’s protocol. T cells were activated for 24 hours with Dynabeads^®^ Mouse T-Activator CD3/CD28 (Life Technologies, Grand Island, NY) in the presence of human IL-2 (30 U/ml) and murine IL-7 (10 ng/ml; PeproTech) in RPMI 1640 media supplemented with 10% heat-inactivated FBS, 1 mmol/L sodium pyruvate, 2 mmol/L glutamine, 100 U/mL penicillin, and 100 μg/mL streptomycin at 37°C with 5% CO2. Activated cells were then transduced by culturing them for 48h in retronectin-coated plates loaded with supernatant harvested from the stable producer line 48h after culture. Invariant NKT cell purity was evaluated by flow cytometry using PE-conjugated PBS-57-loaded mCD1d tetramer (NIH Tetramer Facility) and TCR-β (clone H57-597; BioLegend). Transduction efficacy was measured by flow cytometry after protein L staining (25). Cell numbers were adjusted based on transduction efficacy (50% on average) before *in vitro* or *in vivo* use.

### *In vitro* cytotoxic assay

Murine CD19.28z CAR iNKT cells were co-cultured with luciferase-transduced A20 cells *(A20^yfp+/Iuc+^)(26)* at different ratios adjusted based on transduction efficacy in culture medium consisting of RPMI 1640, supplemented with L-glutamine (2 mM), penicillin (100 U/mL), streptomycin (0.1 mg/mL), 2-mercaptoethanol (5x 10^-5^ M), and 10% FBS. After 24 hours of culture, D-luciferin (PerkinElmer) was added at 5 μg/ml and incubated for 5 min before imaging using an IVIS Spectrum imaging system (Perkin Elmer). Data were analyzed with Living Image Software 4.1 (Perkin Elmer).

### *In vivo* bioluminescence imaging

For *in vivo* bioluminescence imaging (BLI), mice were injected with D-luciferin (10 mg/kg; intraperitoneally) and anesthetized with isoflurane. Imaging was conducted using an IVIS Spectrum imaging system (Perkin Elmer) and data were analyzed with Living Image Software 4.1 (Perkin Elmer) or using an Ami LED-illumination based imaging system (Spectral Instruments Imaging, Tucson, AZ) and data analyzed with Aura Software (Spectral Instruments Imaging).

### *In vivo* murine tumor models

We employed two systemic B-cell lymphoma mouse models previously reported (26). Briefly, CD19-expressing BCL_1_^*luc+*^ (5 × 10^4^) or *A20^luc+^* cells (2 × 10^4^) resuspended in PBS were injected intravenously (i.v.) by tail vein into alymphoid BALB/c (H-2K^d^) Rag1^-/-^ gamma-chain^-/-^ mice. For tumor induction in immunocompetent mice, tumor cells were injected i.v. into sublethally (4.4 Gy) irradiated BALB/c mice. For syngeneic bone marrow transplantation, BALB/c mice were lethally irradiated (8.8 Gy in 2 doses administered 4 hours apart) and transplanted with syngeneic BALB/c bone marrow cells (5 × 10^6^) after T-cell depletion using CD4 and CD8 MicroBeads (Miltenyi Biotec). For retransfer experiments, bone marrow cells from alymphoid BALB/c (H-2K^d^) Rag1^-/-^ gamma-chain^-/-^ mice were used to exclude any potential contribution from bone-marrow derived T or NK cells after reconstitution.

### Flow cytometry analysis

*In vitro* cultured cells or *ex vivo* isolated cells were resuspended in phosphate-buffered saline (PBS) containing 2% FBS. Extracellular staining was preceded by incubation with purified FC blocking reagent (Miltenyi Biotech). Cells were stained with: TIM3 (clone: RMT3-23) APC, CD62L (clone: MEL-14) AF700, CD19 (clone: 6D5) APCFire750, CD44 (clone: IM7) PerCpCy5.5, PD-1 (clone: 29F.1A12) BV605, CD8a (clone: 53-6.7) BV650, NK1.1 (clone: PK136) BV711, ICOS (clone: C398.4A) BV785, CD25 (clone: PC61.5) PE, TCRβ (clone: H57-597) PE/Dazzle594, and Thy1.1 (clone: HIS51) PeCy7. All antibodies were purchased from Biolegend. Dead cells were excluded using Fixable Viability Dye eFluor^®^ 506 (eBioscience). Samples were acquired on a BD LSR II flow cytometer (BD Biosciences), and analysis was performed with FlowJo 10.5.0 software (Tree Star).

### RNA and TCR sequencing analysis

Host CD4 and CD8 T cells were FACS-sorted from pooled spleens from 3 mice treated with allogeneic CAR iNKT or untreated control, frozen in TRIzol and conserved at −80°C. RNA was extracted using the TRIzol RNA isolation method (ThermoFisher Scientific) combined with the RNeasy MinElute Cleanup (Qiagen). Full-length cDNA was generated using the Clontech SMARTer v4 kit (Takara Bio USA, Inc., Mountain View, CA) prior to library generation with the Nextera XT DNA Library Prep kit (Illumina, Inc., San Diego, CA). Libraries were pooled for sequencing on the Illumina HiSeq 4000 platform (75 bp, paired-end). Sequencing reads were checked using FastQC v.0.11.7. Estimated transcript counts and transcripts per million (TPM) for the mouse genome assembly GRCm38 (mm10) were obtained using the pseudo-aligner Kallisto. Transcript-level abundance was quantified and summarized into gene level using the tximport R package. Differential gene expression was performed using the DESeq2 R package version 1.22.221, using FDR < 0.05. Gene-set enrichment analysis conducted using the fgsea R package. For TCR sequencing, libraries were prepared from the synthesized full-length cDNA using the nested PCR method previously reported (28, 29). Sequencing was performed by using the Illumina MiSeq platform after Illumina paired-end adapters incorporation. TCRβ sequence analysis was performed with VDJFasta. After total count normalisation, downstream analysis was performed on the 1000 most represented clonotypes across the samples using the FactoMineR and factoextra R packages.

### Statistical analysis

The Mann-Whitney U test was used in cross-sectional analyses to determine statistical significance. Survival curves were represented with the Kaplan-Meier method and compared by log-rank test. Statistical analyses were performed using Prism 8 (GraphPad Software, La Jolla, CA) and R version 3.5.1 (Comprehensive R Archive Network (CRAN) project (http://cran.us.r-project.org) with R studio version 1.1.453.

## Results

### Allogeneic CAR iNKT cell antitumor effect is significantly enhanced in the presence of host lymphocytes

To study the interaction of allogeneic CD19-specific CAR iNKT cells with the host immune system, we utilized a fully murine experimental system and transduced murine iNKT cells expanded *ex vivo* from FVB/N mice with a previously reported CAR construct (23) composed of the variable region cloned from the 1D3 hybridoma recognizing murine CD19 linked to a portion of the murine CD28 molecule and to the cytoplasmic region of the murine CD3-ζ molecule (CD19.28z CAR; Figure 1A). The cytotoxic potential of CD19.28z-CAR iNKT was confirmed by *in vitro* cytotoxic assays against the CD19-expressing A20 lymphoma cell line, revealing dose dependent cytotoxicity of the CD19.28z-CAR iNKT cells (Figure 1B). As predicted, untransduced iNKT did not display any significant cytotoxic effect against A20 cells (Figure 1B) according to their lack of expression of CD1d. We next evaluated *in vivo* the direct antitumor effect of allogeneic CAR iNKT cells using BALB/c (H-2K^d^) Rag1^-/-^ gamma-chain^-/-^ mice as recipients (Figure 1C, F). FVB/N (H-2K^q^) derived allogeneic CAR iNKT cells significantly controlled tumor growth (Figure 1D) and improved animal survival (Figure 1E) compared to both untreated mice and mice receiving untransduced iNKT cells after administration to major histocompatibility complex (MHC)-mismatched immunodeficient mice receiving CD19-expressing BCL_1_ B cell lymphoma cells. In a second, more aggressive model of B-cell lymphoma using A20 cells (Figure 1F), allogeneic CAR iNKT minimally affected tumor growth as revealed by BLI (Figure 1G) and slightly but significantly improved survival (Figure 1H) compared to untreated mice and mice treated with untransduced iNKT cells.

**Figure 1.**
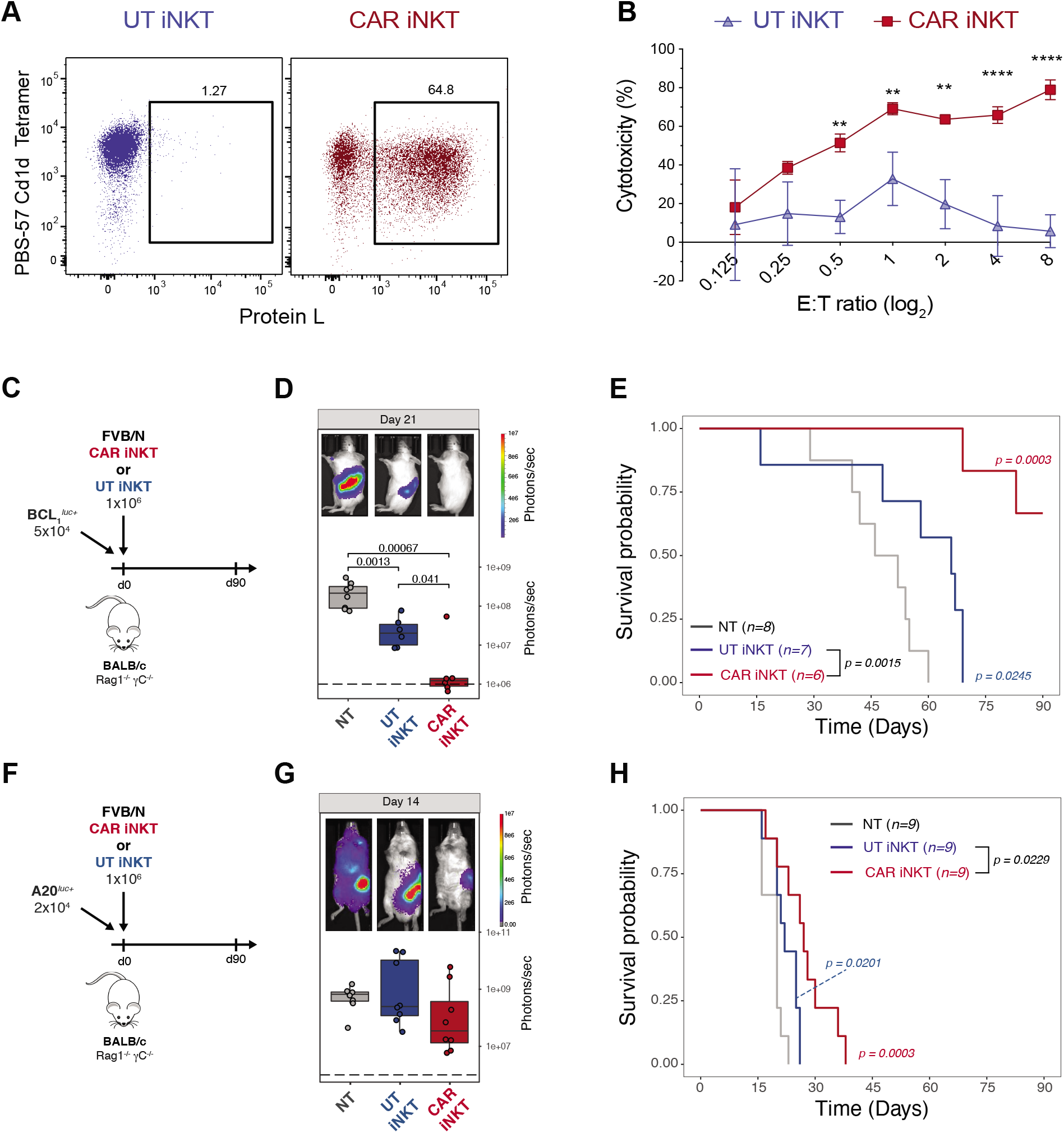
*In vitro* and *in vivo* antitumor activity of murine CAR iNKT cells. (A) Representative FACS-plot of untransduced (left panel) and mCD19.28z-transduced (right panel) murine CAR iNKT cells. iNKT were identified as PBS-57 CD1d tetramer positive cells and CAR transduction was quantified by Protein L staining. (B) Mean and SD of cytotoxicity relative to the untreated control at different E:T ratios. Results are representative of two independent experiments performed in triplicate. (C,F) Schematic representation of the BCL_1_^*luc+*^ (C) *and A20^luc+^* (F) into Rag1^-/-^ gamma-chain^-/-^ recipients experiments. (D,G) Representative *in vivo* bioluminescence (BLI) images of BCL_1_^*luc+*^ (D) *and* A20^*luc+*^ (G) tumor cell progression in Rag1^-/-^ gamma-chain^-/-^ treated with untransduced iNKT cells (blue boxes and dots), CAR iNKT cells (red boxes and dots) or untreated (grey box and dots). (E,H) Survival of mice receiving BCL_1_^*luc+*^ (E) or A20^*luc+*^ (H) and treated with untransduced iNKT cells (blue lines), CAR iNKT cells (red lines) or left untreated (NT, grey lines). Results are pooled from two independent experiments with a total of 6-9 mice per group. BLI results were compared using a nonparametric Mann-Whitney U test and p values are shown when significant. Survival curves were plotted using the Kaplan-Meier method and compared by log-rank test. P values are indicated when significant.

To assess the interplay between the transferred CAR iNKT cells and the host immune cells, we employed the A20 tumor model to test the antitumor activity mediated by allogeneic CAR iNKT cells in an immunocompetent model (Figure 2A) using as recipients wild-type BALB/c mice receiving sublethal irradiation (4.4 Gy) leading to a partial and transient lymphopenia. The antitumor effect of 1×10^6^ allogeneic untransduced iNKT and CAR iNKT cells was greatly enhanced in this partially lymphopenic model, leading to long-term survival of all treated mice (Figure 2B). Interestingly, a dose as low as 5×10^4^ untransduced iNKT cells (Figure 2C) was sufficient to significantly extend animal survival (Figure 2D) and the addition of the CAR further improved the effect of iNKT leading to long-term survival of all CAR iNKT treated mice (Figure 2D). To further stress the model, we tested the antitumor effect of untransduced iNKT and CAR iNKT cells in a high-burden, pre-established tumor model in which high numbers (2.5 x 10^5^) of A20 cells were injected 7 days before the adoptive of the effector cells (Figure 2F). In this model, untransduced iNKT displayed a minimal although statistically significant effect (Figure 2F) while the administration of CAR iNKT cells significantly improved animal survival compared to both untreated mice and mice receiving untransduced iNKT cells (Figure 2F).

**Figure 2.**
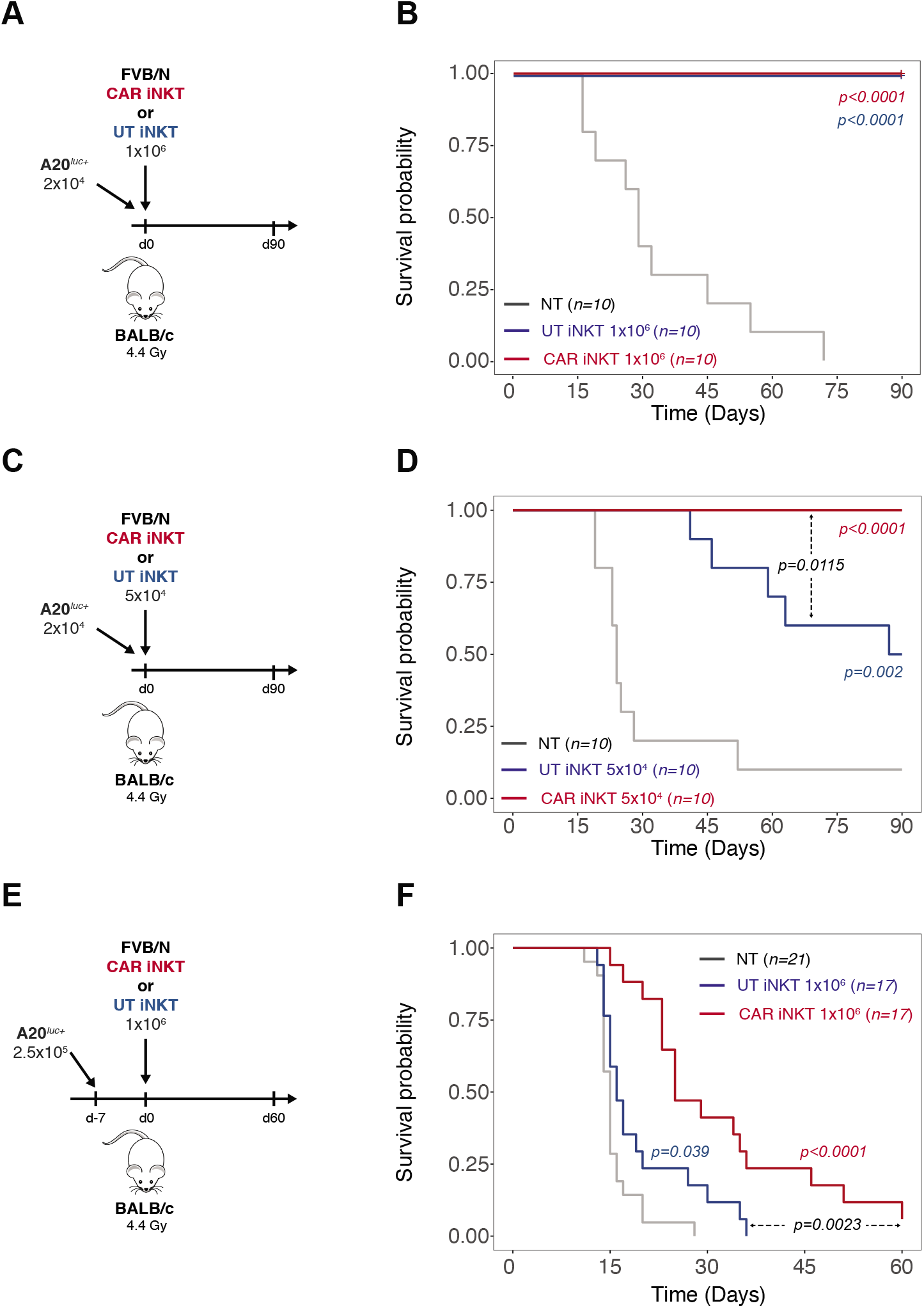
Allogeneic CAR iNKT cell antitumor effect is greatly enhanced by the presence of host lymphocytes. (A,C,E) Schematic representation of the experiments employing *A20^luc+^* cells into sublethally (4.4 Gy) irradiated WT BALB/c mice. (B,D,F) Survival of mice receiving *A20^luc+^* cells and treated with untransduced iNKT cells (blue lines), CAR iNKT cells (red lines) or left untreated (NT, grey lines). Results are pooled from two independent experiments with a total of 10-21 mice per group. Survival curves were plotted using the Kaplan-Meier method and compared by log-rank test. P values are indicated when significant.

Collectively, these *in vitro* and *in vivo* data confirm the direct antitumor effect of murine CAR iNKT cells and revealed an improved effect of untransduced iNKT and, even more, of CAR iNKT cells in the presence of host lymphocytes.

### Host CD8 T cell cross-priming contributes to the indirect antitumor effect of allogenic CAR iNKT cells

The striking difference in allogeneic CAR iNKT effect observed in mice with partial lymphopenia (Figure 2B, D) compared to genetically alymphoid mice (Figure 1H) suggested a role for host-derived lymphocytes in the antitumor effect. To test the hypothesis that host CD8 T cell cross-priming mediates the indirect antitumor effect of allogenic CAR iNKT cells, we employed as recipients BALB/c BATF3^-/-^ mice, in which CD8 T cell cross-priming is impaired as a result of the absence of BATF3-dependent CD103+ CD8alpha+ dendritic cells (30). The effect of allogeneic CAR iNKT cells was partially abrogated in A20-receiving BATF3^-/-^ mice as compared to WT mice (Figure 3A-B), supporting the hypothesis that the impact of allogeneic CAR iNKT cells is mediated, at least partially, by the activation of host CD8 T cells via their cross-priming. To further assess the synergistic effect of allogeneic CAR iNKT cells and host-derived CD8 T cells, we employed an autologous bone marrow transplantation model, co-administering allogeneic FVB/N CAR iNKT with syngeneic BALB/c CD8 T cells at the time of transplantation with T-cell-depleted syngeneic BALB/c bone marrow cells and transfer of A20 lymphoma cells into lethally irradiated (8.8 Gy) BALB/c recipients. Co-administration of allogeneic CAR iNKT and autologous CD8 T cells resulted in a synergistic effect, significantly improving tumor control (Figure 3C) and animal survival (Figure 3D) compared to mice receiving no treatment, as well as to mice receiving either allogeneic CAR iNKT or autologous CD8 T cells alone. Collectively these data indicate that CD8 T cell cross-priming is necessary for allogeneic CAR iNKT cells to exert their full antitumor effect and suggest a synergy between these two cytotoxic T cell compartments.

**Figure 3.**
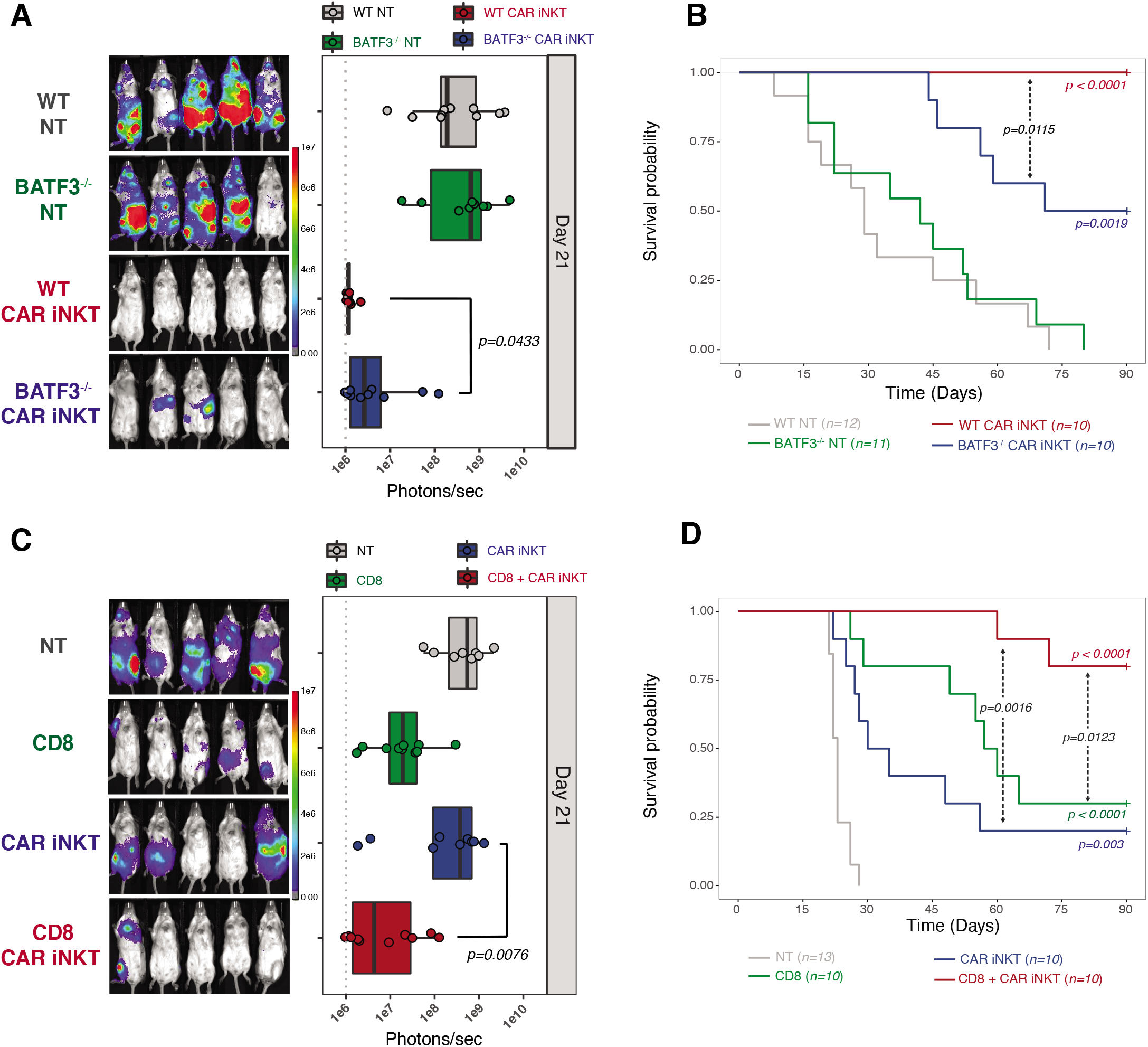
Indirect antitumor effect of allogeneic CAR iNKT cells is dependent on host CD8 T cells cross-priming. Representative *in vivo* BLI images of A20^*luc+*^ cell progression (A) and survival (B) of sublethally (4.4 Gy) irradiated WT or BATF3^-/-^ BALB/c mice treated or not with 10^6^ CAR iNKT cells. Representative *in vivo* BLI images of *A20^luc+^* cell progression (C) and survival (D) of lethally (8.8 Gy) irradiated WT BALB/c mice transplanted with syngeneic BALB/c TCD-BM and treated with syngeneic CD8 T cells (4×10^6^; green symbols and line), CAR iNKT cells (10^6^; blue symbols and line) or both (red symbols and line). Untreated controls are depicted in grey. BLI results were compared using a nonparametric Mann-Whitney U test and p values are shown. Survival curves were plotted using the Kaplan-Meier method and compared by log-rank test. P values are indicated when significant.

### Allogeneic CAR iNKT cell treatment modulates host CD8 T cells phenotype, transcriptome and TCR repertoire

To gain further insights into the impact of CAR iNKT cells on host T cells, we performed phenotypic analysis of host T cells recovered at day 7 and 14 after treatment with allogeneic CAR iNKT cells. At these timepoints, allogeneic CAR iNKT cells had been already rejected as revealed by *in vivo* tracking by bioluminescence (Supplemental Figure 1A), flow cytometry (data not shown) and as suggested by the progressive increase of B cell numbers (Supplemental Figure 1B). We observed a significant increase in the number of CD8 T cells recovered at day 7 and day 14 from the spleen of mice treated with CAR iNKT cells compared with untreated mice (Figure 4A). Immunophenotypic analysis revealed higher proportions of cells with a central memory (CD62L+ CD44+) and reduced proportions of cells with an effector (CD62L-CD44-) or effector memory (CD62L-CD44+) phenotype in CD8 T cells recovered at day 7 after allogeneic CAR iNKT treatment compared with untreated mice (Figure 4B). CD4 T cell numbers were increased at day 7 but not at day 14 after allogeneic CAR iNKT treatment (Supplemental Figure 2A), and CD4 T cell phenotype was only minimally affected by CAR iNKT treatment (Supplemental Figure 2B). A transcriptomic analysis performed on CD8 T cells FACS-sorted at day 14 revealed the upregulation of genes associated with cytotoxic antitumor activity (*Lyz2, Gzma, Gzmm, Fasl*) and the downregulation of genes involved with responses to type I interferon (*Irf7, Ifitm1, Ifi27l2a;* Figure 4C). Gene Set Enrichment Analysis (GSEA) for Gene Ontology (GO) Biological Processes confirmed the upregulation of antitumor gene sets (Figure 4D) and the downregulation of the type I interferon signature. In agreement with our phenotypic results, a GSEA performed using two well-established memory CD8 T cell gene signatures revealed enrichment in CD8 T cell memory genes (Figure 4E-F). Transcriptomic analysis of CD4 T cells showed a similar downregulation of genes involved in responses to type I interferon (*Irf7, Ifit1, Ifit3;* Supplemental Figure 2C) but did not reveal any consistent pattern of expression of genes involved in antitumor activity or cellular differentiation (Supplemental Figure 2C). To assess the impact of CAR iNKT treatment on the TCR repertoire of CD8 T cells, we performed paired TCR-beta sequencing. Hierarchical clustering based on the 1000 most represented TCR clonotypes revealed a closer relationship between the TCR repertoire of CD8 T cells recovered from mice receiving CAR iNKT cells compared with CD8 T cells from untreated mice (Figure 4G). Accordingly, principal component analysis (PCA) showed close similarity in the TCR repertoire of CD8 T cells from allogeneic CAR iNKT treated mice, while cells from untreated mice displayed high heterogeneity (Figure 4H). Analysis of the TCR repertoire of CD4 T cells did not reveal any impact of allogeneic CAR iNKT treatment (Supplemental Fig. 2D). Collectively, these results indicate that allogeneic CAR iNKT cell treatment shaped the host CD8 T cell compartment phenotypically, transcriptomically, and in terms of clonal repertoire.

**Figure 4.**
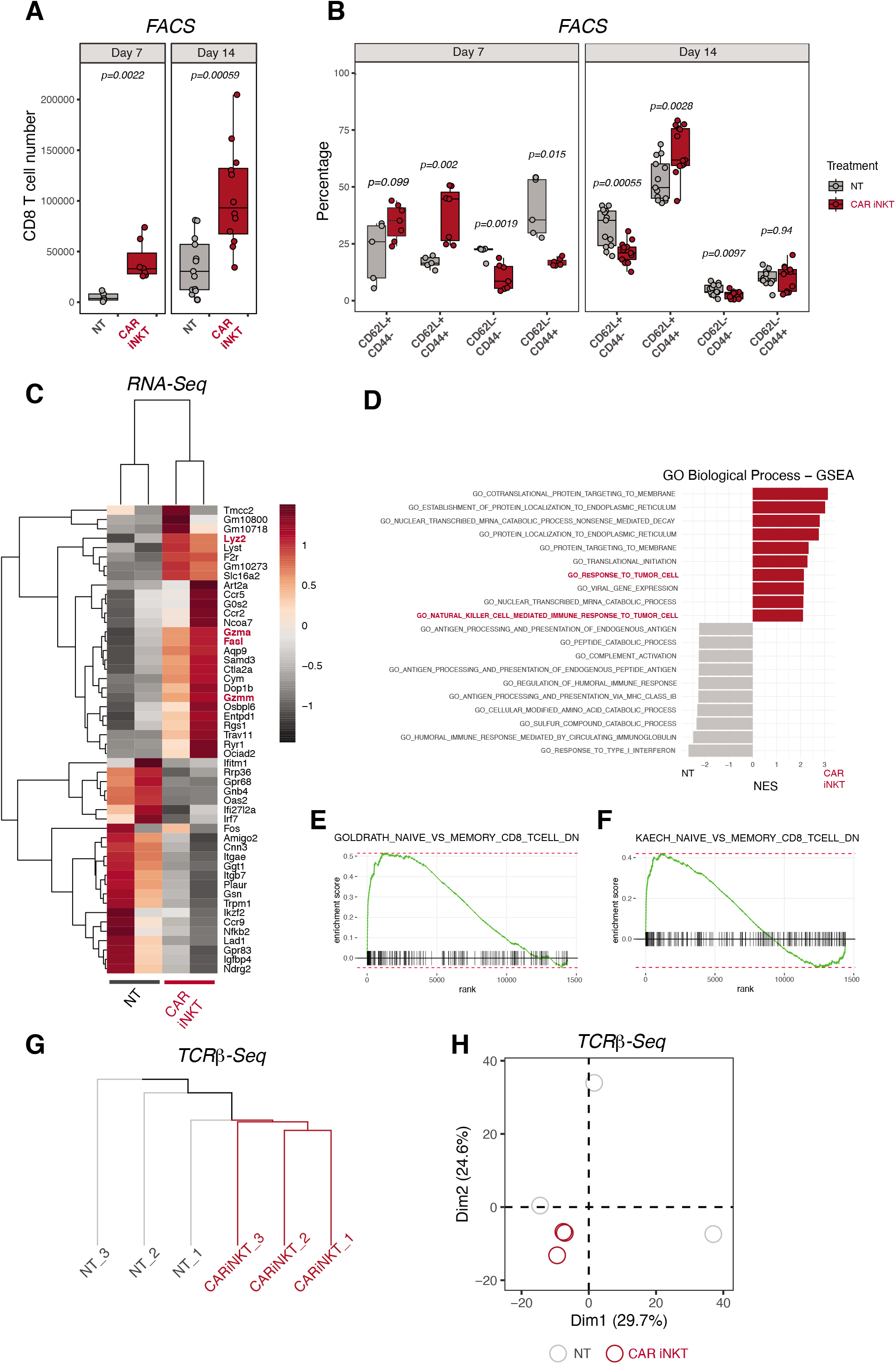
Allogeneic CAR iNKT cell treatment modulates host CD8 T cell number, phenotype, transcriptome, and TCR repertoire. Number (A) and immunophenotype (B) of host CD8 T cells recovered from spleen 7 and 14 days after tumor induction in mice treated with allogeneic CAR iNKT cells (red boxes and symbols) or untreated (grey boxes and symbols). Results are pooled from two independent experiments with a total of 5-13 mice per group. Groups were compared using a nonparametric Mann-Whitney U test and p values are shown. (C) Heatmap representing differentially expressed genes in host CD8 T cells FACS-sorted from recipients treated or not with allogeneic CAR iNKT cells. Expression for each gene is scaled (z-scored) across single rows. Each column represents independent experiments with one to two biological replicates per experiment. (D) Top 10 enriched terms/pathways in CD8 T cells from untreated (grey bars) and CAR iNKT cell treated (red bars) animals revealed by GO Biological Process analysis using GSEA. (E-F) Enrichment plots displaying the distribution of the enrichment scores for the genes down-regulated during transition from naive CD8 T cells versus memory CD8 T cells according to the *Goldrath et al.* (E) or *Kaech et al.* (F) signatures. Gene signatures were obtained from Molecular Signatures Database (MSigDB; C7: immunologic signatures). (G) Hierarchical clustering and (H) principal component analysis (PCA) of the top 1000 clonotypes based on TCRß sequencing of CD8 T cells from hosts treated with allogeneic CAR iNKT cells (red) or left untreated (grey).

### Allogeneic CAR iNKT cell treatment induces long-lasting host CD8 T cell tumor-specific responses

To formally prove that allogeneic CAR iNKT cells induce tumor-specific host immune responses, at day 60 after treatment we recovered splenocytes from mice receiving A20 lymphoma cells and treated with allogeneic CAR iNKT cells. Recovered splenocytes were transferred into new lethally irradiated BALB/c recipients together with bone marrow from Rag1^-/-^ gamma-chain^-/-^ BALB/c mice and A20 cells (Figure 5A). Unprimed splenocytes from mice receiving only sublethal irradiation were used as control. Splenocytes primed in the presence of allogeneic CAR iNKT significantly extended the survival of mice compared to both untreated mice and mice receiving unprimed splenocytes (Figure 5B, left panel). To assess the contribution of CD8 T cells to this protective effect, we performed the same experiment retransferring only allogeneic CAR iNKT-primed or unprimed CD8 T cells. As shown in Figure 5B (right panel), host CD8 T cells from allogeneic CAR iNKT-treated mice significantly extended animal survival compared to both mice left untreated or receiving unprimed CD8 T cells. Collectively, these experiments formally demonstrate that allogeneic CAR iNKT treatment induced a long-lasting tumor-specific host CD8-dependent antitumor immunity in allogeneic recipients.

**Figure 5.**
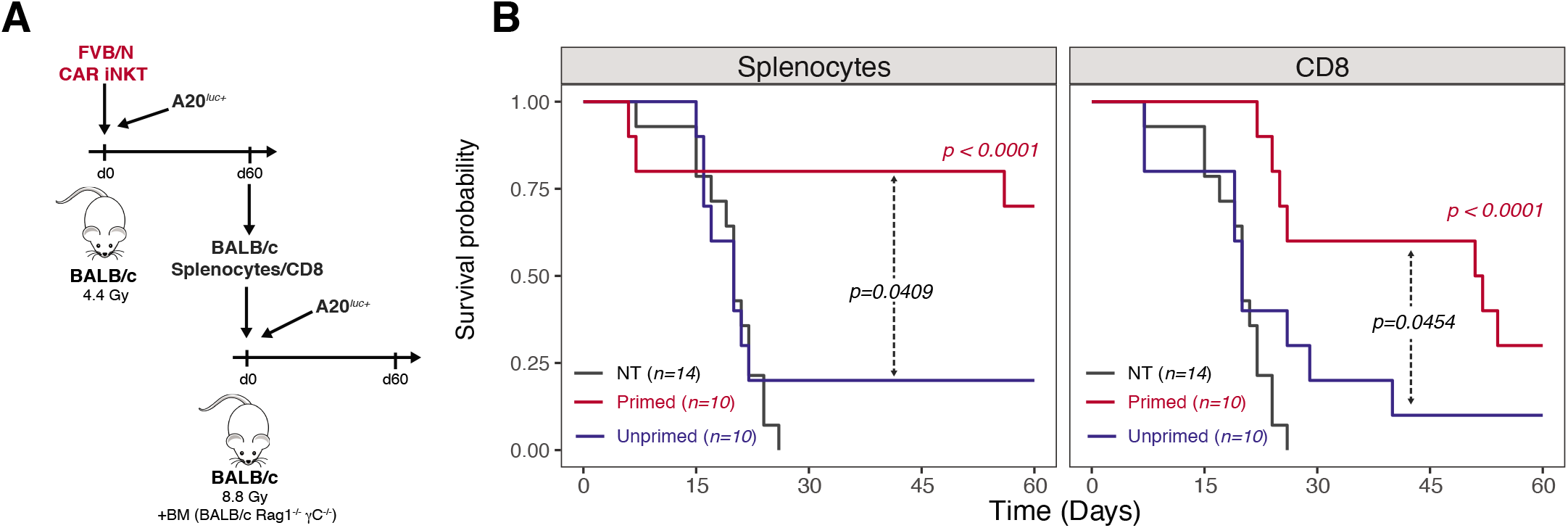
Allogeneic CAR iNKT cell-primed host CD8 T cells display long-lasting antitumor immunity. (A) Schematic representation of the sequential adoptive transfer experiment. Host splenocytes or CD8 T cells were recovered after 60 days from sublethally irradiated BALB/c mice, injected with A20^*luc+*^ cells, and treated with CAR iNKT cells (primed cells). Splenocytes or CD8 T cells recovered after 60 days from sublethally irradiated BALB/c mice were used as controls (unprimed cells). Primed or unprimed host splenocytes (5×10^6^ cells) were transferred, after lethal irradiation, to a new set of BALB/c mice receiving *A20^luc+^* cells together with bone marrow cells from syngeneic Rag1^-/-^ gamma-chain^-/-^ BALB/c mice. Alternatively, primed or unprimed host CD8 T cells (1×10^6^ cells) were transferred. (B) Survival of transplanted mice receiving primed (red line) or unprimed (blue line) splenocytes (left panel) or CD8 T cells (right panel). Untreated controls are depicted in grey. Results are pooled from two independent experiments with a total of 10-14 mice per group. Survival curves were plotted using the Kaplan-Meier method and compared by log-rank test. P values are indicated when significant.

### Allogeneic CAR iNKT cells outperform conventional CAR T cells in the presence of host lymphocytes

To assess the advantage that this indirect antitumor effect could confer to allogeneic CAR iNKT cells over allogeneic conventional CAR T cells, we compared these two populations. Given the potent direct antitumor activity of conventional CAR T cells, a dose of as little as 2.5×10^5^ conventional CAR T cells was sufficient to significantly extend mouse survival when administered into alymphoid animals (Supplemental Figure 3), and this dose was selected for comparison to CAR iNKT cells. As shown in Figure 6A-B, during partial lymphopenia conventional CAR T cells significantly extended animal survival, while CAR iNKT cells dramatically outperformed conventional CAR T cells leading to tumor control and survival of all treated mice. Collectively, these results demonstrate that allogeneic CAR iNKT cells were significantly more effective than allogeneic conventional CAR T cells in inducing extended tumor control in immunocompetent hosts.

**Figure 6.**
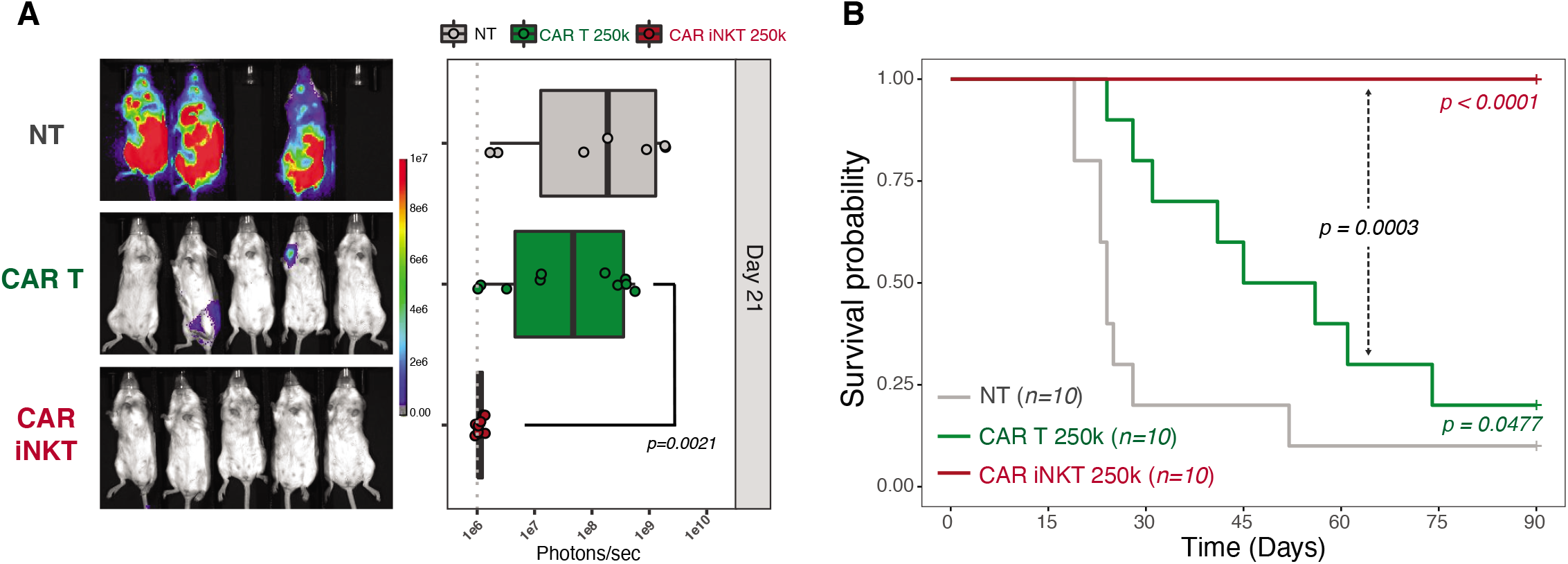
Allogeneic CAR iNKT cells are more effective than allogeneic conventional CAR T cells. Representative *in vivo* BLI images of *A20^luc+^* cells progression (A) and survival (B) of sublethally (4.4 Gy) irradiated BALB/c mice treated with 2.5×10^5^ allogeneic CAR iNKT (red curve and symbols), 2.5×10^5^ allogeneic conventional CAR T (green curve and symbols) cells or untreated (grey curve and symbols). Results are pooled from two independent experiments with a total of 10 mice per group. BLI results were compared using a nonparametric Mann-Whitney U test and p values are shown. Survival curves were plotted using the Kaplan-Meier method and compared by log-rank test. P values are indicated when significant.

## Discussion

In the present study, we demonstrated in a murine model of CD19+ lymphoma that allogeneic CAR iNKT cells exert, in addition to their previously reported direct antitumor effect, an even stronger indirect antitumor effect mediated by the induction of host immunity.

The potential contribution of the host immune system in the effect of CAR T cells has been shown in preclinical (31–35) and clinical (36, 37) studies. In particular, Beatty et al. showed that transiently expressed mRNA CARs were able to induce epitope spreading despite their limited persistence (36). We hypothesized that induction of bystander antitumor responses might be a particularly interesting approach in the allogeneic setting, as CAR cells administered across major-MHC barriers will be invariably rejected by the host immune system. After confirming in a fully murine model the previously reported direct cytotoxic effect of CAR iNKT cells (8–12), we demonstrate that allogeneic CAR iNKT cells efficiently induce anti-tumor immune responses in the recipient through host CD8 T cell cross-priming. Importantly, our phenotypic and transcriptomic data indicate that host CD8 T cells primed in the presence of allogeneic CAR iNKT display a central memory profile and we provide evidence that retransfer of CAR iNKT primed CD8 T cells allow for the transfer of protective antitumor immunity. These results suggest that the antitumor effect lasts much longer than the physical persistence of the administered allogeneic cells.

iNKT cells are an ideal platform for off-the-shelf immunotherapies given their lack of GvHD-induction potential (38) without need for deletion of their endogenous TCR, a manipulation that has been recently shown to alter the CAR T cell homeostasis and persistence (39). Moreover, despite being a rare lymphocyte population, iNKT cells can be easily expanded *ex vivo* to numbers needed for clinical uses (40–43) and several clinical trials using *ex vivo* expanded autologous iNKT cells have been already successfully conducted (44–46). However, previous reports indicate that the ability of iNKT cells to expand *in vitro* may vary widely among individuals(47), a potential limitation for generation of autologous or allogeneic MHC-matched products. Use of allogeneic, off-the-shelf iNKT cells to be administered across MHC barriers will circumvent this potential limitation as universal donors whose iNKT cells display optimal expansion potential can be selected. Moreover, our results indicate that extremely low numbers of CAR iNKT cells persisting for a very limited time are able to induce a potent, long-lasting antitumor effect through their immunomodulatory role. Such an effect is in accordance with what we previously reported in the GvHD settings, where similarly low numbers (5×10e4) of CD4+ iNKT cells were able to efficiently prevent GvHD induced by conventional T cells in a major MHC-mismatch mouse model of bone marrow transplantation (48), even when rapidly rejected third-party cells were employed (49).

A phase I clinical trial employing CD19-specific allogeneic CAR iNKT cells for patients with relapsed or refractory B-cell malignancies is currently ongoing (ANCHOR; NCT03774654). In analogy to what performed with conventional CAR T cell, this clinical trial involves the administration of a lymphodepleting regimen containing fludarabine and cyclophosphamide before CAR iNKT cell infusion. Our results indicate that a major component of allogeneic CAR iNKT cells effect derives from their interplay with the host immune system, an interaction that can significantly be impaired by the lymphodepleting conditioning. Future studies will determine whether the conventional fludarabine/cyclophosphamide lymphodepletion interferes with CAR iNKT cell effect and will test alternative regimens to optimize both the homeostasis and the immunoadjuvant effect of the administered product.

In conclusion, our results represent the first demonstration of an immunoadjuvant effect exerted by an allogeneic CAR cell product toward the host immune system, resulting in long-lasting antitumor effects that go beyond the physical persistence of the allogeneic cells.

## Author contributions

FS conceived and designed research studies, developed methodology, conducted experiments, acquired and analyzed data, and wrote the manuscript; JKL, TH, KMB, MA, ASW conducted experiments; XJ, developed methodology and analyzed data; JB, AA, SH developed methodology and provided essential reagents; CLM provided essential reagents and intellectual input; RSN provided overall guidance and wrote the manuscript.

## Data and materials availability

Sequencing datasets will be made publicly available upon acceptance and prior to final publication.

## Supplemental Figure Legends

**Supplemental Figure 1. Limited persistence and absence of B-cell aplasia after allogeneic CAR iNKT cell treatment.** (A) Persistence of *Luc+* CAR iNKT cells in tumor bearing mice at different time points. BLI data are expressed as photon/sec in mice receiving 1×10^6^ *Luc+* CAR iNKT cells after subtracting the background detected in mice not receiving Luc+ cells. Data shown are from two merged independent experiments with 5 mice per group in each experiment. (B) B cell numbers. B cells were defined by FACS as CD19+ YFP- to distinguish normal B cells from CD19+ YFP+ A20 cells. Median (black dashed line) and upper/lower range (gray dotted lines) of B cell counts in naive mice are represented.

**Supplemental Figure 2. Limited impact of allogeneic CAR iNKT cell treatment on host CD4 T cells.** Number (A) and immunophenotype (B) of host CD4 T cells recovered from spleen 7 and 14 days after tumor induction in mice treated with allogeneic CAR iNKT cells (red boxes and symbols) or untreated (grey boxes and symbols). Results are pooled from two independent experiments with a total of 5-13 mice per group. Groups were compared using a nonparametric Mann–Whitney U test and p values are shown. (C) Heatmap representing differentially expressed genes in host CD4 T cells FACS-sorted from recipients treated or not with allogeneic CAR iNKT cells. Expression for each gene is scaled (z-scored) across single rows. (D) Principal component analysis (PCA) of the top 1000 clonotypes based on TCRß sequencing of CD4 T cells from hosts treated with allogeneic CAR iNKT cells (red) or left untreated (grey).

**Supplemental Figure 3. Direct antitumor effect of conventional CAR T cells in alymphoid mice.** Survival of alymphoid BALB/c Rag1-/- gamma-chain-/- mice receiving 2.5×10^5^ (solid green lines), 5×10^4^ (dashed green lines) allogeneic conventional CAR T cells or untreated (NT, grey lines). Results are pooled from two independent experiments with a total of 6-8 mice per group. Survival curves were plotted using the Kaplan-Meier method and compared by log-rank test.

