## Supplementary figures and images for "Allogeneic CAR-invariant Natural Killer T Cells Exert Potent Antitumor Effects Through Host CD8 T Cell Cross-Priming"

### Supplemental Figure 1

#### SUPPLEMENTAL FIGURE 1

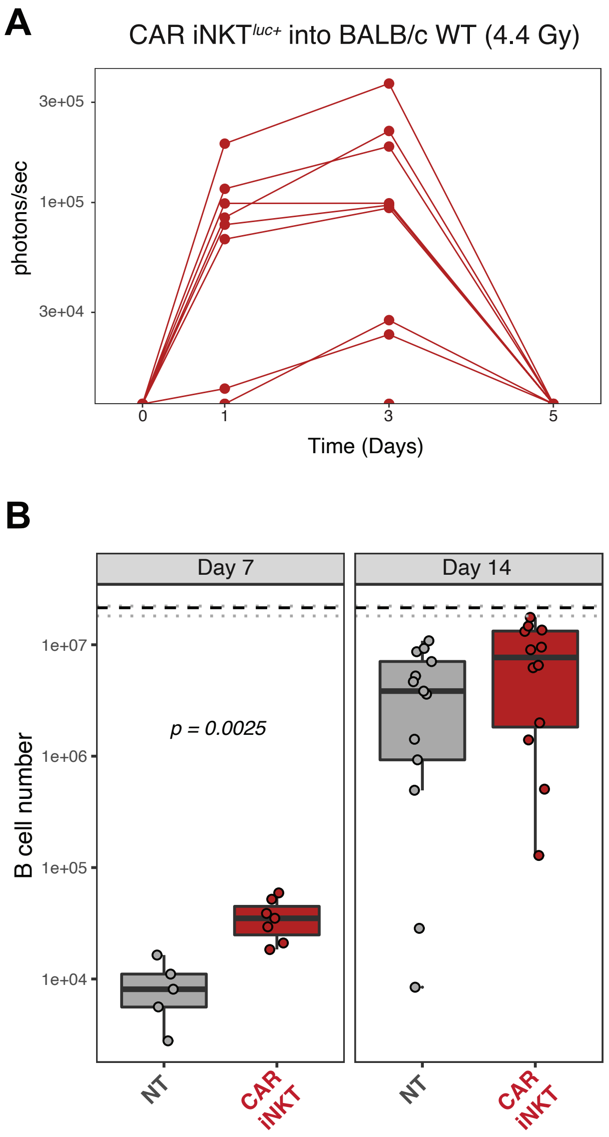

### Supplemental Figure 2

#### SUPPLEMENTAL FIGURE 2

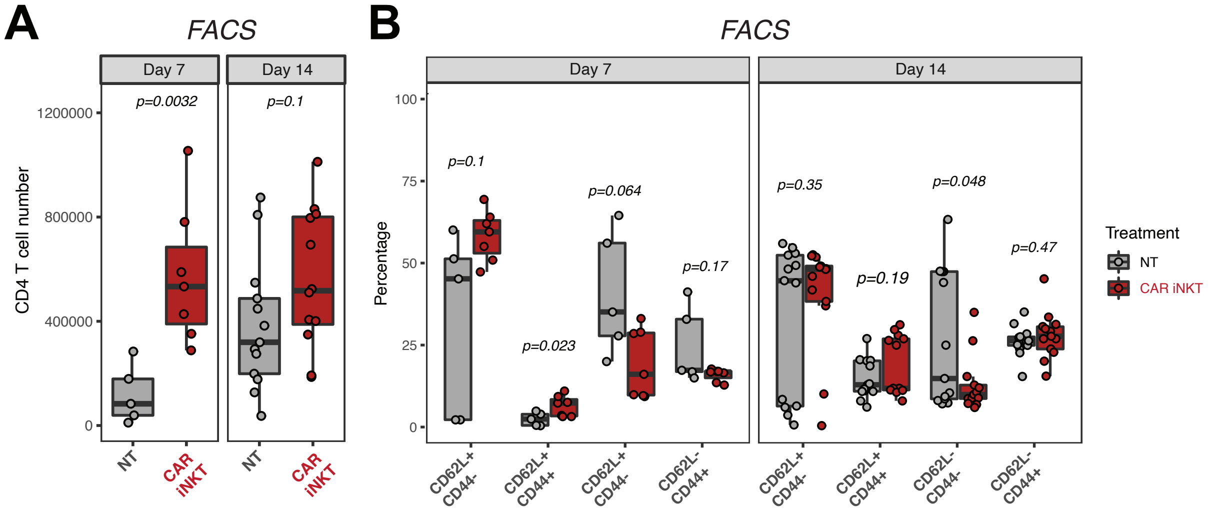


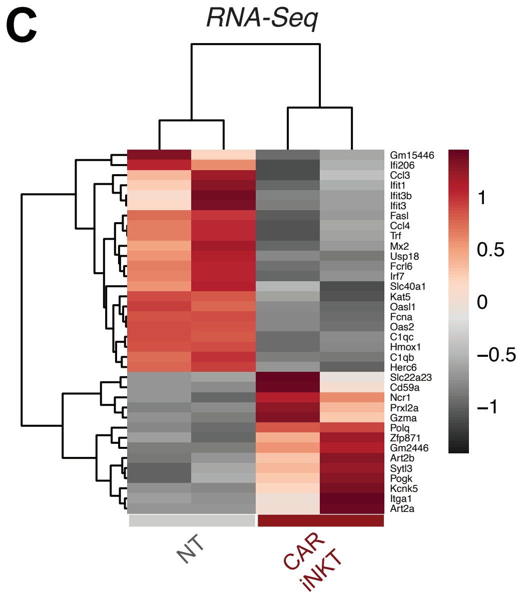


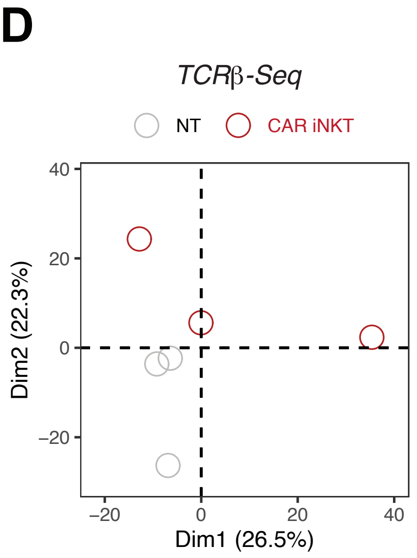

### Supplemental Figure 3

#### SUPPLEMENTAL FIGURE 3

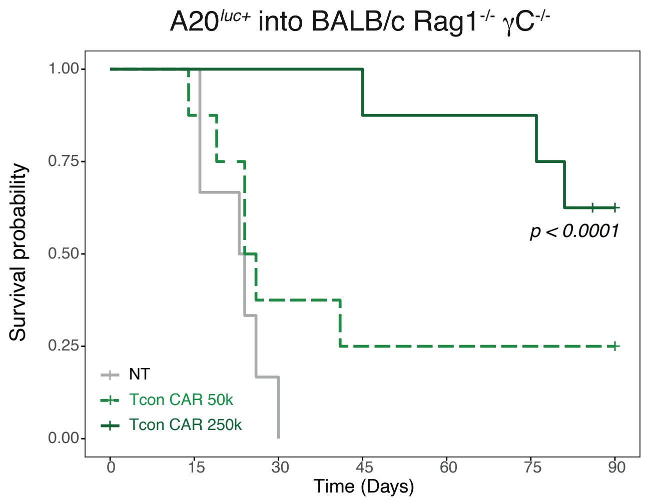
